# Influence of the ecological opportunity of interaction on the structure of host-parasite networks

**DOI:** 10.1101/2020.01.13.904151

**Authors:** Elvira D’Bastiani, Karla Magalhães Campião, Walter Antonio Boeger, Sabrina Borges Lino Araújo

## Abstract

Despite the great interest to quantify the structure of ecological networks, the influence of morphological, ecological and evolutionary characteristics of the species still remains poorly understood. One of the challenging issues in ecology is how the interaction opportunity influences and provides changes to the associations between species, and which effects these changes have on ecological systems. To explore topological patterns in host-parasite networks, we sampled endoparasites-anurans interactions in South America in order to determine whether the effect of the ecological opportunity affects our understanding of the topological structure of the interaction networks. To identify the effect of the ecological opportunity for interaction, we investigated interactions in environments with and without flood pulse, where presence would promote higher ecological opportunity of interaction. Moreover, we created three theoretical models with filters to test the influence of the ecological opportunity for interaction: random, phylogeny and host body size. We then calculated commonly used binary network metrics (connectance, nestedness and modularity) for the networks generated by the theoretical models. We demonstrated that the interaction ecological opportunity changes the structure of host-parasite networks, and was influenced mainly by phylogeny and body size of the host. Our results indicate that environments that offer greater opportunities for interaction between species present networks with the most connectance/nestedness and less modularity. Networks in environments that do not have such opportunities for interaction depict the opposite pattern. Our results indicate that the ecological opportunity of interaction is reflected in an increase in interaction associations between species and affect/change the organization of these interactive assemblages. From an epidemiological point of view, changes in the composition of parasitic species are associated with risks of invasions and emerging diseases. In part, emerging diseases are the result of processes such as those occurring during the flood pulse, in which climate change, travel, and global trade create opportunities for new species associations. Our results provide insight into the dynamics of incorporating a new resource, considering an evolutionary factor responsible for these changes in species composition.

## Introduction

Understanding factors that determine the establishment and persistence of interactions between species is fundamental to comprehend the factors that structure ecological communities [40]. Host-parasite interactions are influenced by characteristics of the hosts and associated parasites. For hosts, characteristics such as body size and structure, diet, age, and immune system are considered significant [43,48,23], but these and others are the resources that dictate the feasibility of the association with specific parasite species [7, 15]. Important characteristics on the parasite species include life cycle complexity [23], virulence, transmissibility, strategies to evade immune responses [65], host species recognition, and trophic requirements [45], among others. Therefore, the effects of these features directly influence the structure of interaction networks.

Hence, intrinsic factors/characteristics of each association determine the potential for an interaction to occur, and compose what Combes [21], Araujo et al. [7], and [15] designate as compatibility. However, the effective establishment of a compatible interaction depends on the encounter opportunity [7,15,73]. This effectiveness is often the result of ecological adaptation [38], which postulates that key characteristics shared with previous host species are necessary for a successful infection [33,12,74]. The ecological adaptation is the process that allows the parasite organism to successfully colonize a new host without the need of evolutionary novelties [3,50].

The support to this new vision of parasitism lies in the fact that the macroevolutionary process, commonly named host-switching, has proven to be much more common in nature than previously thought [74,15]. Many human pathogens originate through host-switching, including HIV and malaria [76], and host-shifts are also the predominant cause of new host-parasite associations for rabies viruses in bats [70], malaria in birds [28], and among other associated diseases [14, 50, 55,59].

The ecological opportunity for interaction is significantly influenced by the environment [15]. Cyclical changes in expansion and isolation, for instance, generate Taxon Pulse processes [27, see also 34]. Taking this into account, Taxon Pulse processes can lead to increase or limitation of opportunities for interaction between hosts and parasites and, consequently, alter the structure of the community. The flood pulse, for example, is considered a key factor in the ecological functioning and the patterns of lowland communities [63,71]. These flood pulses [41] tend to reduce spatial variability and biological as well as environmental factors [9,71]. According to this hypothesis, during low water periods, floodplains are more isolated from each other and disconnected from the main channel of the river, creating isolated habitats, often with distinct environmental characteristics. On the other hand, subsequent increases in water levels represent an expansion event for aquatic and semiaquatic organisms. We postulate that this pattern of isolation/expansion between neighboring aquatic habitats (flood plain/flood pulse system) provides the ecological opportunity for encounter between endoparasites and anurans species, which may result in new interactions through ecological adjustment. By contrast, continuously isolated habitats provide more limited opportunities for encounter between hosts and parasitic species. In fact, some studies have reported greater similarity between species in the composition of different aquatic habitats during floods than during non-flood periods [5,9,63,72]. We also expect that the structure of communities subjected to cycles/pulses of isolation and expansion will present greater sharing and connectivity of endoparasites between hosts. Limited ecological opportunity in continuously isolated habitats should exhibit less parasite sharing and lower endoparasites connectivity among the anuran hosts, changing the structure of the network.

In this study, we tested whether this structure is influenced by phylogeny and body size hosts and whether the periodic ecological opportunity of encounters influences the structure of an anuran-endoparasite (endoparasite-anuran) interaction network in environments with and without annual flood pulses. We used a database of anuran-endoparasite interactions to infer how the network structure of each environment can be explained by randomly sampled networks obtained under different theoretical models with filters: neutral, host phylogeny, and host body size. If phylogeny or body size imposes restrictions on species interactions, we expect to find compartments in the random networks. On the other hand, if there are no restrictions, networks would show nested patterns with interaction indirectly reflecting the abundance of species. We conclude that, among the filters analyzed, host body size and phylogeny greatly explain the observed host-endoparasite networks. However, similar networks of environments with cyclic flood pulse were not well explained by these filters, strongly suggesting that the increased opportunities provided by the environment can intensify encounters and promote and increase connectivity in the host-parasite local network.

## Materials and Methods

To test for differences in ecological opportunity, we selected four parasite-host interaction networks compiled from literature data and described them using network metrics (nestedness, connectance, and modularity). Next, we sought to know if it was possible to explain their structure based on the phylogenetic characteristics and body size of the hosts analyzed. For this, we created theoretical models with filters (see below) and then compared the structure of the four networks analyzed with the networks generated by theoretical models to test the effect of the process of interaction of opportunity in different environments. See the methods for details.

### Host-parasite database

We created a database with reports on the association of interactions between anurans and helminth parasites (endoparasite) from South America. All possible combinations (e.g. amphibians, endoparasites, helminth, anura) were used to search for anuran-endoparasite (endoparasites belonging to the phylum Acanthocephala, Nematoda, and Platyhelminthes, associated with amphibians of the order Anura) empirical studies conducted in South America from 01-Jan-1925 to 20-Dez-2017. These data were collected using online database platforms such as BioOne, Isi JSTOR, PubMed, SciELO, Scopus, and Web of Science. We have updated the amphibian’s nomenclature according to the American Museum of Natural History [29]. *Leptodactylus latrans* interaction reports were not included in the analyses due to many changes in nomenclature. From here on we will call "host" for anuran and "parasite" for helminth parasites. We then generated a binary matrix with these real interactions in which the rows represented host species and the columns represented parasite species. This matrix was used as an interaction database to generate random networks through the theoretical models (filters). We collected 157 peer-reviewed articles and recorded 686 real interactions between 215 species of parasites and 170 species of hosts (Appendix S1).

### Selection of analyzed environments

During the bibliographical review, four (4) community studies of host-parasite interactions (observed networks) were selected: (i) two studies of flood pulse environments (with annual cycles of water expansion and retraction), which could promote environmental homogenization and higher ecological opportunity for host-switching: Pantanal - Campião et al. [18], and Chaco - González and Inés [30] flood pulse. The Pantanal network was composed by 11 hosts and 16 parasites (Appendix S2) while the Chaco, by 35 and 46, respectively (Appendix S2) [18,30]. (ii) Two studies from environments without flood pulse, which potentially promote less ecological opportunity for host-switching: Atlantic Rainforest Graça et. [31], and Amazonian forest Bursey et al. [16]. The Atlantic Rainforest network was composed by 11 host and 15 parasites (Appendix S2), while the Amazonian forest by 43 and 16 respectively (Appendix S2) [31,16].

### Theoretical models

To answer our questions, we chose to test the effect of random, phylogeny, and host body size to see the changing structure (ecological opportunity) of host-parasite networks in different environments. We chose phylogeny because it was revealed as a potential driver of parasitic diversity. Host species vary in their evolutionary time of exposure for parasites acquisition and sharing, therefore, suffering variable co-evolutionary constraints [32] which would influence the interactions and the structure of their networks’ interaction. We also chose the host body size as it was a good predictor of the diversity of parasite species [17,42,53]. Large hosts can provide more space and resources, and possibly a greater breadth of niches for parasites. Moreover, larger hosts live longer and represent fewer ephemeral habitats than small species and are therefore also more exposed to parasites [61].

Three theoretical models with filters were created (random, phylogeny and body size of hosts models - Appendix S3), to test the influence of the ecological opportunity on the structure of host-parasite networks. Each model is characterized as a specific filter that randomly selects hosts and their parasites from the interaction database that resulted in random networks. Unlike what is commonly used to analyze the structure of networks, our random networks considered only real interactions extracted from the interaction database (collected from the literature). Our models have generated random networks using real interactions and may give a more accurate answer about the topology of the interaction network than other simpler null models commonly used that consider only random interaction simulations, or that weigh only by the interaction ratio of the species network for example. Given this approach the following theoretical models were proposed: i. Neutral Model: No other filter besides the number of host species, being the same number in each observed network (11 hosts for Pantanal, 35 for Chaco, 11 for Atlantic Rainforest and 43 for Amazonian forest respectively - Appendix S3 - Filter i); ii. Phylogeny Model: This model randomly samples from the database the same number of host species belonging to the same families in the same proportion of the original network, as reported in each observed network (Appendix S3 - Filter ii; Appendix S4 - families of the species by observed network); iii. Body Size Model: This model randomly samples from the database the same number of hosts species with the same body size distribution (considering a standard error of ±5%) as reported for hosts species in each observed network (Appendix S3 - Filter iii; Appendix S4 - body size of the species by observed network).

For each one of the four environments analyzed, the random networks maintained the number of host species of each observed network (Pantanal, Chaco, Atlantic Rainforest and Amazonian forest). In this way, the number of hosts remained constant (according to each environment), while the number of parasites varied between the random networks, according to the association in the database of recorded interaction.

We sampled 1.000 random networks for each model in each environment (random networks per model in Appendix S5). Each random sample was always included in the interactions database after each randomization, so that a species could be included in more than one random network. We then compared how well the filters explain the observed network (the comparison methods are detailed below). Unfortunately, it was not possible to impose on a similar real filter the ecological opportunity to evaluate the effect of interaction opportunity because we have no observation that refers to such a factor. Aquatic and terrestrial systems have different physical and chemical conditions, which seem to influence the biology and diversity of the organisms living in each of these habitats [75], that is why we assumed phylogenetic conservatism of the organisms living in each of these analyzed habitats. Given their aspects, we hypothesized that flood pulse environments are not as well explained by the filters (neutral, phylogeny, and host body size) as the other environments. We postulate that in environments without flood pulse the phylogenetic conservatism could be higher, that is, as it is an environment with few opportunities the species maintain their interactions over evolutionary time. If this is true, we expect that the networks will be better described by the phylogeny and body size models that are supposed to have their most preserved interactions. This is something we would not observe in the cyclic flood pulse because there is a constant change that interferes with this conservatism.

### Data analysis

#### Network Metrics

We used the metrics nestedness, connectance, and modularity to characterize the structure of all networks. The nestedness was calculated using the metric proposed by Almeida-Neto et al. [6], to evaluate the presence of interactions that belong to subsets of other interactions. Therefore, a high nestedness value indicates a hierarchy of interactions, in which species that interact with fewer partners (have a lower degree) interact with a subset of partners from species that have more partners (a higher degree) [8]. Connectance is the proportion of interactions performed for all possible interactions between species in a community [52]. Modularity was calculated using the method proposed by Dormann and Strauβ [24], to describe the presence of network groupings, where species interact more with species within their group than with species belonging to other groups [58]. These groups are commonly called network modules, and to calculate these network metrics, we used the commands implemented in the “bipartite” package [25] in R [22].

#### Standardization for network metrics

To allow comparison between networks of different sizes, we standardized the network metrics using a simple linear regression between metrics and number of parasites. This standardization is necessary because although the host species richness was constant in the different samples, the size of the whole network differed in each sample due to variation in parasite richness. See below:

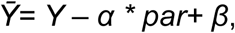

where *Ȳ* is the value of the standardized metric (equivalent to the regression residue), *Y* is the value of the non-standardized metric, *par* is the number of parasites, and *α* and *β* are the slope and the linear coefficient of the regression, respectively.

#### Comparison of the network structure

From here onwards we call "observed network" any of the four observed networks for tested environments, and "random network" any network generated by filters. We identify which model best explains each observed network as well as which network is best described by the models. To compare networks, we first calculated the distance (*D*) for each metric, from each random network of each theoretical model to the observed network in units of standard deviation:

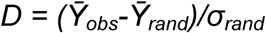

where *Ȳ*_*obs*_ is the standardized value of the observed network, *Ȳ*_*rand*_ is the normalized value of the random network metric, and *σ*_*rand*_ the standard deviation of 1.000 values of *Ȳ*_*rand*_. Subsequently, for a given metric and observed network, a unidirectional analysis of variance (ANOVA) was applied to evaluate whether there was a difference between these groups. Tukey’s test [79] was used to determine the occurrence of differences between treatments. After measuring the difference, the distance averages between the real networks and each random network of each model was compared to verify which theoretical model best described the observed networks. When the metric and the observed network were fixed, we compared which model best explained the observed network. When the metrics and model were fixed, we compared which observed network was best explained by the models. All statistical analyses were performed using the "stats" package in the R software [22]. For all tests, we assumed the significance of *p*<0.05.

## Results

### Structure of host-parasite interaction networks

Some of the random networks presented a range from smaller to greater number of parasite species than the observed networks (Fig. 1, 2, 3 - black bars, upper x axis - frequency of the number of parasites), except for the Amazon, whose smallest random network had at least twice more parasites than the observed network. Thus, by correcting the effect of network size, the theoretical models could not reach a network as small as that analyzed in the Amazon due to low parasitic richness analyzed in this environment (Fig. 1, Fig. 2 and Fig. 3 - j to l). Despite our attempt to control the effect of network size by a standardization method, we were unable to ensure that the ratio of the metric and the number of parasites maintained the same linear tendency for such low numbers of parasites for Amazon. Therefore, the Amazonian simulations were removed from subsequent analysis.

**Fig. 1.**
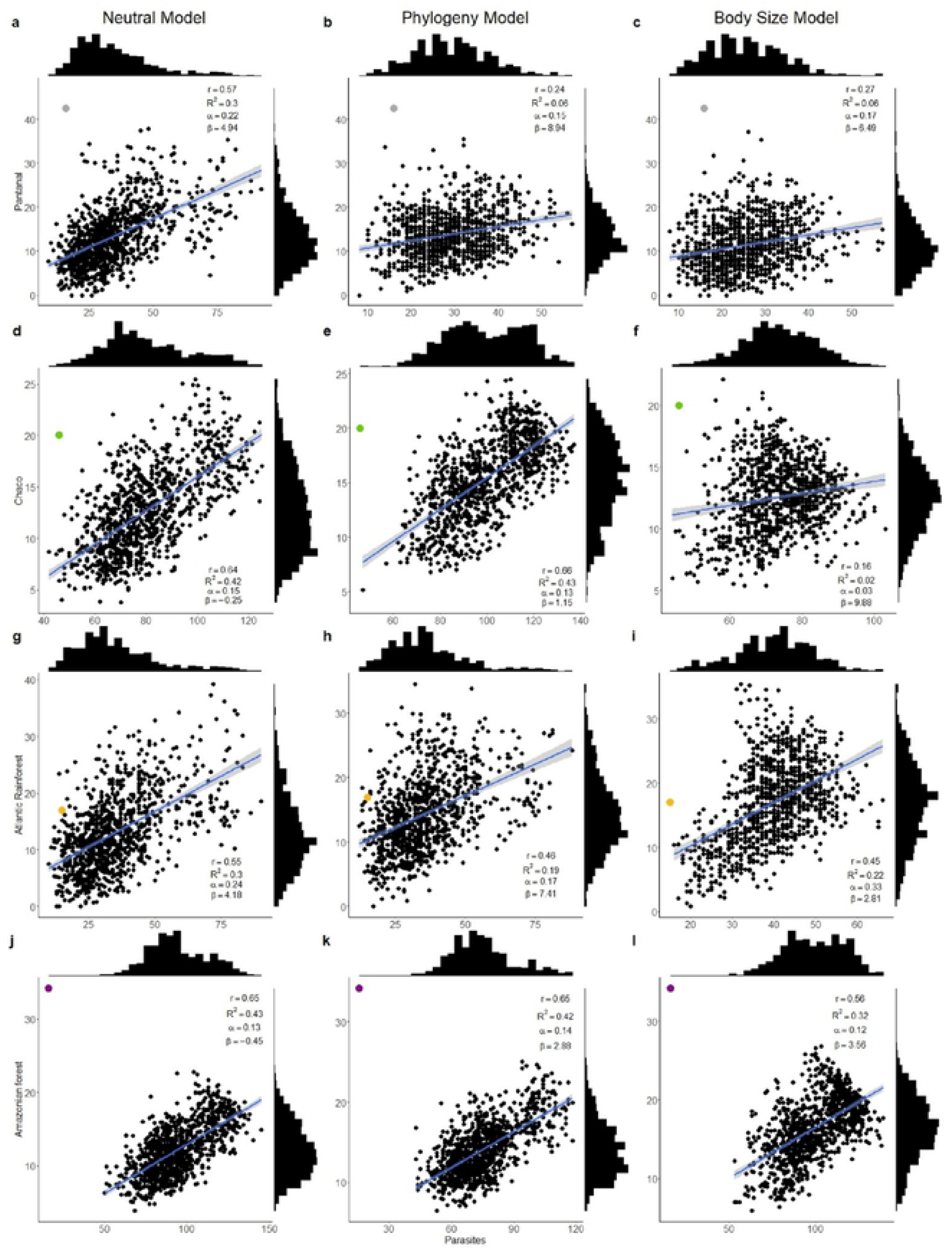
Nestedness values according to the number of parasite species before standardization. The black dots indicate the nestedness value of random networks for three theoretical models. Grey dot - Pantanal (a, b, c), green dot - Chaco (d, e, f), orange dot - Atlantic Forests (g, h, i) and violet dot - Amazonian forest (j, k, l). The blue lines represent linear regression and the gray shadow represents standard deviation in random networks. The black bars indicate the frequency distribution of the random networks: right y axis - nestedness frequency, upper x axis - frequency of the number of parasites (details: Appendix S10 in Supporting Information).

**Fig. 2.**
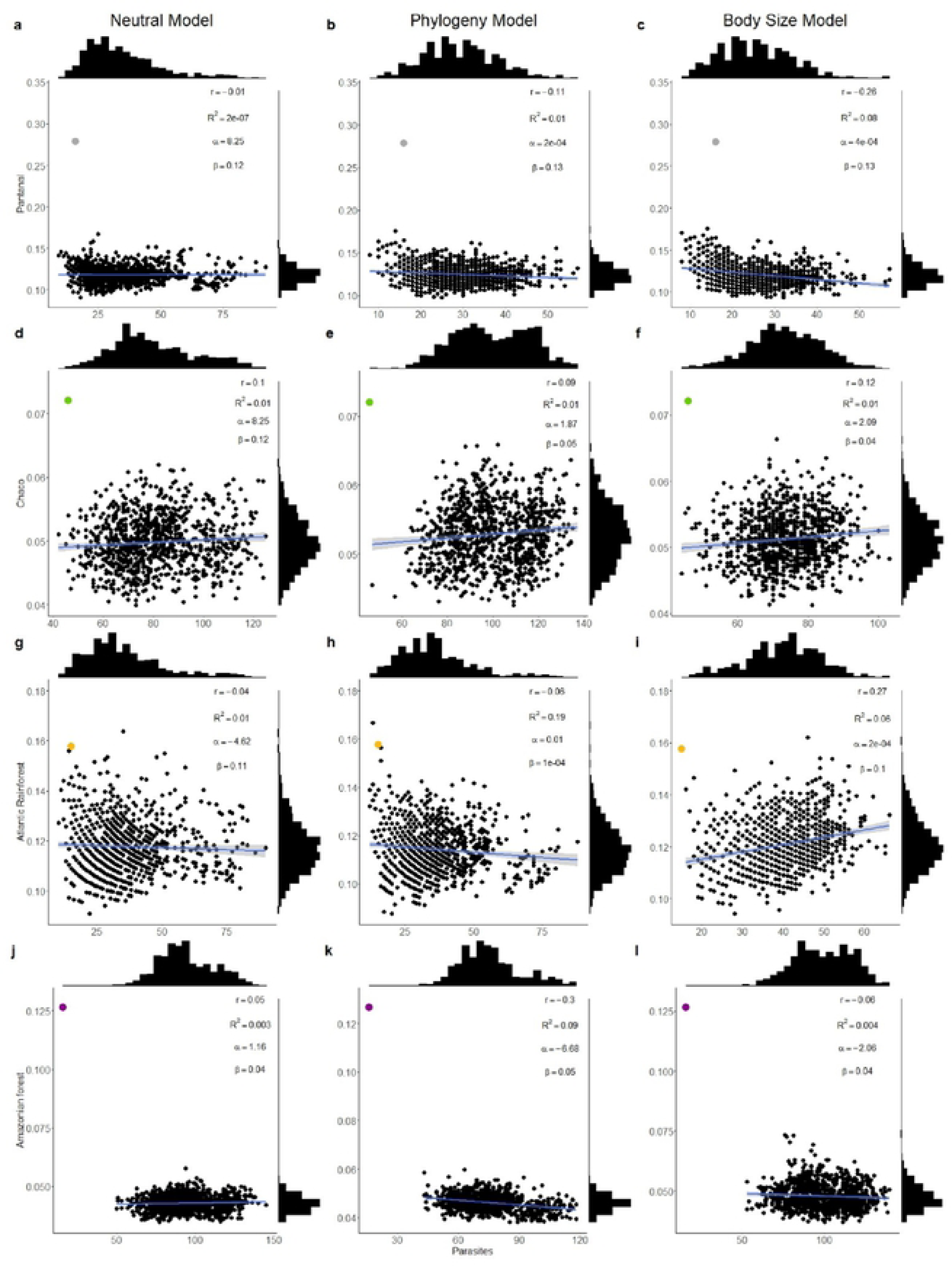
Connectance values according to the number of parasite species before standardization. The black dots indicate the connectance value of random networks for three theoretical models. Grey dot - Pantanal (a, b, c), green dot - Chaco (d, e, f), orange dot - Atlantic Forests (g, h, i) and violet dot - Amazonian forest (j, k, l). The blue lines represent linear regression and the gray shadow represents standard deviation over the random networks. The black bars indicate the frequency distribution of the random networks: right y axis - connectance frequency, upper x axis - frequency of the number of parasites (details: Appendix S10 in Supporting Information).

**Fig. 3.**
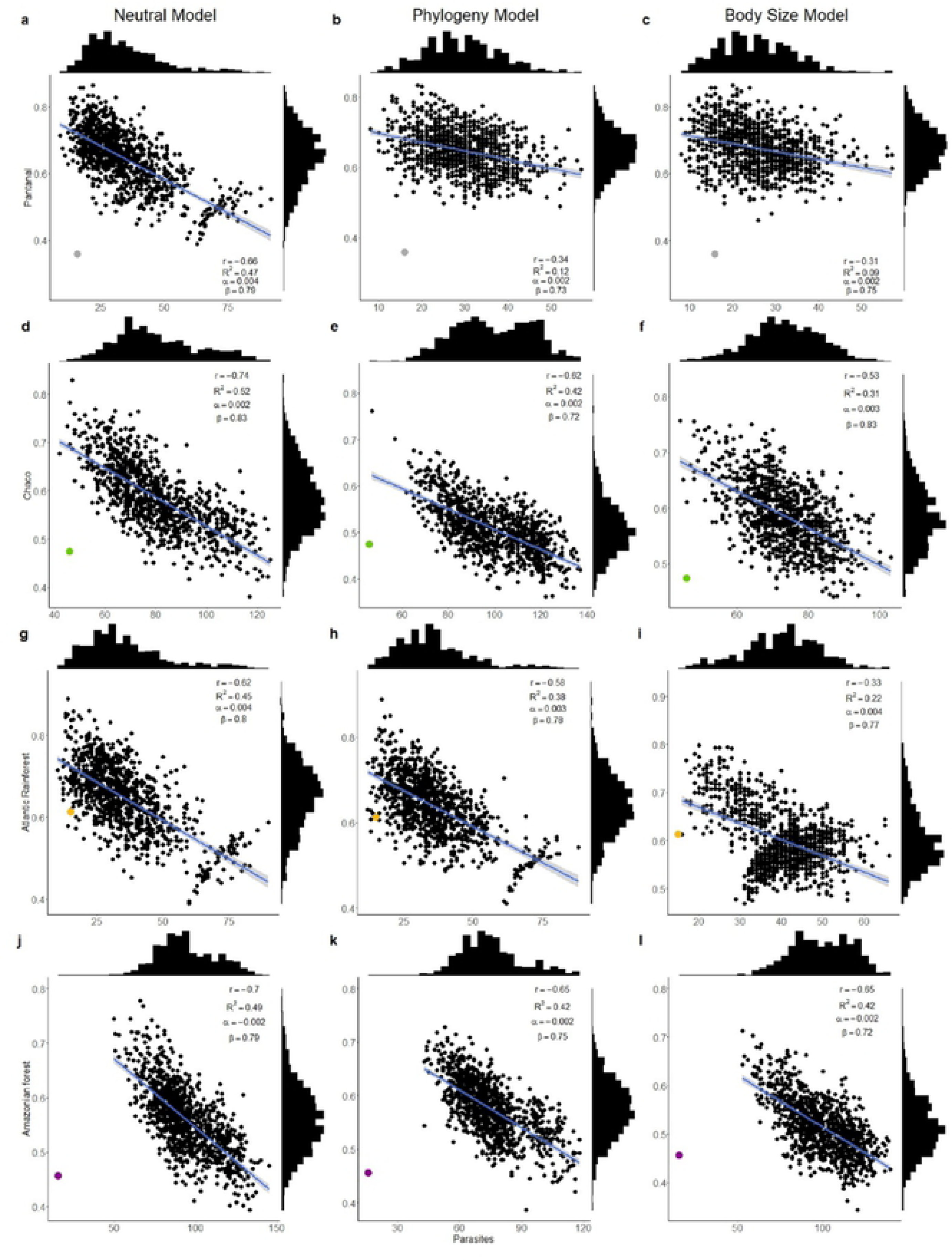
Modularity values according to the number of parasite species before standardization. The black dots indicate the modularity value of random networks for three theoretical models. Grey dot - Pantanal (a, b, c), green dot - Chaco (d, e, f), orange dot - Atlantic Forests (g, h, i) and violet dot - Amazonian forest (j, k, l). The blue lines represent linear regression and the gray shadow represents standard deviation over the random networks. The black bars indicate the frequency distribution of the random networks: right y axis - modularity frequency, upper x axis - frequency of the number of parasites (details: Appendix S10 in Supporting Information).

The nestedness increases with the number of parasites in the network for the remaining three environments (Fig. 1, Appendix S6). Connectance and modularity were negatively correlated with the number of parasites in most simulated models (Fig. 2 and 3, Appendix S6), except for the connectance in simulated neutral models for the Pantanal (R² = 0.01, *p* = 0.98) and Atlantic Rainforest, which were not correlated (R² = 0.002, *p* = 0.15).

### Host body size and phylogeny models usually better describe the observed network than neutral models

The distance (D) between random networks and their respective observed network were significantly different among the models (Table 1). The body size model that better resembled the nestedness pattern was observed in the Pantanal and Chaco, while in the Atlantic Rainforest, it was the phylogeny model. The connectance values in the Atlantic Rainforest were better resembled by the neutral model, while in the Pantanal and Chaco were better resembled by the body size and phylogeny models, respectively. The modularity was better resembled by the phylogeny model in the Pantanal and Chaco networks, while modularity in the Atlantic Rainforest was best explained by the body size model (Table 1). Such results, with the exception for connectance in the Atlantic Rainforest, show that the body size and phylogeny of the hosts resemble better the network structure than neutral model.

**Table 1.**
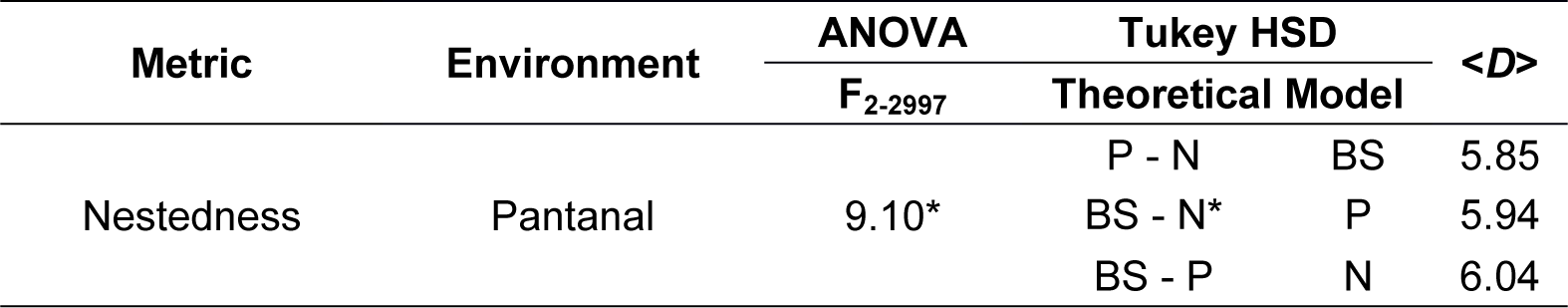
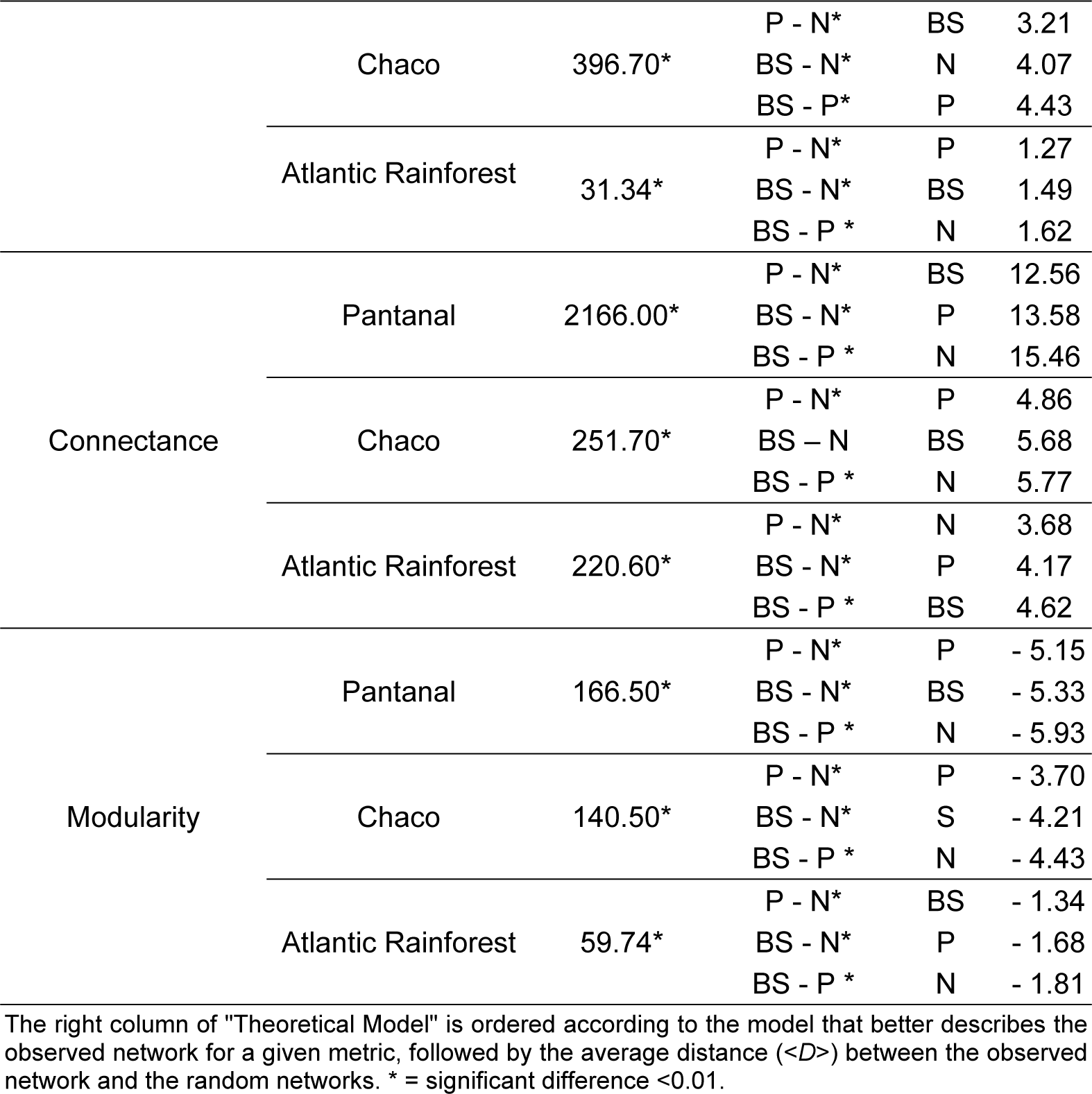
Analysis of variance (statistical significance test F_2-2997_), Tukey Test between neutral models (N), phylogeny (P) and body size (BS) for each metric and environment (observed network).

### The theoretical models describe environments without flood pulse better than the flood pulse environments

The resemblance of the network metrics from randomly sampled networks to observed networks environments with and without flooding for each metric presented different distances (Table 2). All metrics pointed network structure of the Pantanal as the furthest from the network structure of the theoretical models (statistics for support Table 2), followed by the Chaco and the Atlantic Rainforest. This shows that the environment without flood pulse is better described by the proposed models than the flood pulse environments. All theoretical models pointed that Pantanal network is the most nested, connected and less modular one followed by Chaco and Atlantic Rainforest (see the statistics for support test for each metric between random networks of the models and observed network - average distance value, <D>, in Table 2).

**Table 2.**
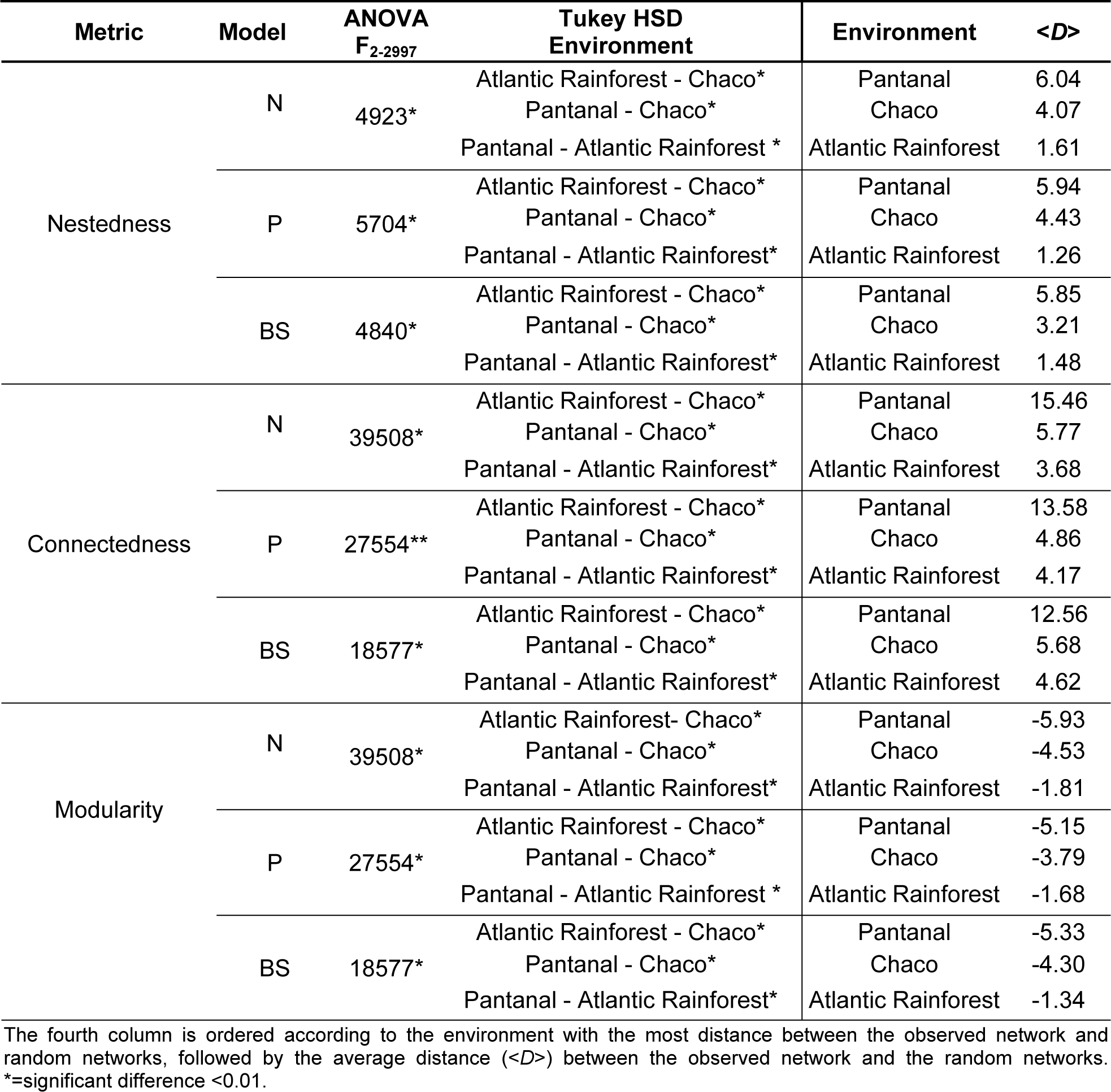
Analysis of variance (statistical significance test F_2-2997_), Tukey Test between neutral (N), phylogeny (P) and body size (BS) theoretical models, for each metric and environment.

## Discussion

The method we developed to infer how the structure of anuran-endoparasite interaction networks under different conditions of encounter opportunities could be described by theoretical models based on neutral, host phylogeny and host body filters. We found that the theoretical models described the network without flood pulse - i.e low ecological opportunity for interaction (Atlantic Rainforest) – better than the networks with cyclic flood pulse (Pantanal and Chaco). In addition, the networks of the three environments were best described by the theoretical models with phylogeny or host body size filters rather than the neutral filter. This result suggests that, in fact, the increased ecological opportunity for encounters provided by the environment increases connectivity through the incorporation of new host species in the repertoire of the parasites.

The topology patterns observed in the communities reflect the mechanisms that structure their respective networks. Some studies suggest randomness as an underlying mechanism for explaining the structure of parasitic communities [44,62]. However, the metrics we tested in our study did not indicate the neutral model as the main structuring mechanism for network topology, except for the connectance of the network in the Atlantic Rainforest. The interactions of the Atlantic Rainforest can be the result of random neutral encounters between individuals, whose probability is mediated by the relative abundances of the populations involved and by compatibility in the case of parasites.

In parasitic communities, the evolutionary history of the host acts as one of the determinants of the community structure [19,67]. Phylogenetic relationships function as a good proxy for describing ecological differences between hosts and host groups. This may indicate that these attributes are phylogenetically conserved, modulating interspecific barriers for parasite colonization among hosts [19,36], and outlining current interactions [26,64].

Many life-history traits are positively correlated with body size and therefore may have affected the structure and dynamics of ecological networks at multiple scales of biological organization, from the individual to the ecosystem [20,60,66,77]. Currently, the major challenge is to develop a body of theory that can explore the implications of body size on the structure and functioning of the host-parasite networks.

The metrics of the Atlantic Rainforest networks were more similar to the theoretical models than the Pantanal and Chaco networks (Fig. 1, 2, 3 and Table 2). These points to other potential factor(s), besides phylogeny and size, acting on the structure of the Pantanal and Chaco networks. Unlike the Atlantic Rainforest, the Pantanal and Chaco are environments marked by large annual floods. As we suggested, the presence of these floods promotes greater contact between aquatic and semi-aquatic species, thus increasing the ecological opportunity for the establishment of new host-parasite associations [2,4]. The encounters can lead to new associations by ecological fitting, i.e. the ability of organisms to adapt quickly to new resources due to their phenotypic flexibility, without genetic novelties [3,7,38].

Moreover, species may expand habitat use in response to the availability of new resources [49,78], which would be associated with the encounter of different hosts in this study. Some examples of habitat expansion (colonization and adaptation to new resources or other new selective ecological adjustment scenarios) arise directly from the events associated with the ecological opportunity [51]. Cyclical expansion of host-parasitic communities in the ecosystems subjected to annual flood pulses favors seasonal contact of infectious forms of parasites with hosts. The colonization of new host species should in fact generate more connected, more nested, and less modular networks (since modules are broken due to the new connections promoted by the environment - Fig. 3, a-f). These events increase the repertoire of the parasite, and may represent the beginning of speciation processes, or, they could maintain its repertoire of host as well as postulated by “Oscillation hypothesis” [10,37,56]. This hypothesis consists largely of micro and macroevolutionary aspects. The microevolutionary part deals with how novel hosts are incorporated during host expansions and, as a consequence, pathogenic lineages can diversify in resource use. The second part is largely macroevolutionary and foresees that these episodes of increased host use should lead to elevated rates of diversification. The particularities of the microevolutionary part affect the specific patterns expected at the macroevolutionary level. In this context, over the course of evolutionary time, true generalist species may become specialists, and vice versa [4,37,56].

In environments without flood pulse, such as the Atlantic Rainforest, the host-parasite interaction may be restricted by the low dispersion of species among aquatic environments, since for many semi-aquatic and aquatic species, the forest environments may represent ecological barriers. As contact is the main route of parasite transmission (expected for endoparasites with heteroxenous and monoxenic cycle), the structure of the Atlantic Rainforest network is very similar to the structure of random networks, since hosts share fewer parasites. Our results suggest that the ecological opportunity for interaction may play an important role in determining the structure of interaction networks and the evolutionary dynamics of host-parasite associations. These results create new perspectives for studies on parasitic community assemblages, particularly as few studies relate the connectivity of host communities to the opportunity for parasitic dispersion. This may, in fact, increase our understanding on the influence of the dynamics of the physical environment on the structure of host-parasites interactions.

From an epidemiological perspective, changes in the composition of parasitic species or in the frequency of host-parasite interactions are associated with the risks of parasitic invasions and emerging diseases [1,4,11,13,33,39,46,54]. Some examples come from studies with the introductions of species [35,57]. In birds, for instance, the occurrence of nematode parasites increases with the ecological opportunity due to migratory habits and the use of aquatic habitat [47,57]. This results in a change in community structure and the formation of new species associations through a combination of mechanisms, such as ecological adaptation and opportunity for interaction between species [68,69]. In part, the emerging diseases are a result of processes such as those that occur during the flood pulse, in which climate change, travel, and global trade generate opportunities for new species associations [57]. We show that an insight is provided into the dynamics of the incorporation of a new resource, as an evolutionary factor considered to be responsible for changes in species composition.

Our results may have been influenced by the fact that the host-parasites interaction database was built from several studies, with different sampling efforts in different regions. This could have increased the interaction records, and, consequently, increased the connectance and reduced the modularity of theoretical models. On the other hand, the interaction database had a larger spatial scale than the local studies in the analyzed environments, so that the random networks could have selected hosts that did not co-occur and, therefore, did not share the same species of parasites, resulting in more modular, but less connected random networks. Even so, the variation in network size between the simulations is high; there is a risk that methodological artifacts (low sample effort) may have been affected, especially in the Amazon network. The fact that the Amazon has a low parasitic richness could be due to the lower sampling effort. Therefore, we opted for excluding this environment from our analyses. However, even with these factors, all the observed networks had the same tendency; they were all more connected and less modular than predicted by the models, thus validating our results.

## Acknowledgements

The authors would like to thank the Academic Publishing Advisory Center (Centro de Assessoria de Publicação Acadêmica, CAPA – www.capa.ufpr.br) of the Federal University of Paraná for assistance with English editing. EDB was supported by a Master degree fellowship from Coordenação de Aperfeiçoamento de Pessoal de Nível Superior (CAPES).

## Author Contributions

KMC, WAB, SBLA and EDB originally formulated the idea. SBLA and EDB developed the mathematical models, conducted work and generated data analyses. EDB, SBLA, KMC and WAB wrote the manuscript.

